# Fine scale population structure and extensive gene flow within an Eastern Nearctic snake complex (*Pituophis melanoleucus*)

**DOI:** 10.1101/2021.09.28.462151

**Authors:** Zachary L. Nikolakis, Richard W. Orton, Brian I. Crother

## Abstract

Understanding the processes and mechanisms that promote lineage divergence is a central goal in evolutionary biology. For instance, studies investigating the spatial distribution of genomic variation often highlight biogeographic barriers underpinning geographic isolation, as well as patterns of isolation by environment and isolation by distance that can also lead to lineage divergence. However, the patterns and processes that shape genomic variation and drive lineage divergence may be taxa-specific, even across closely related taxa co-occurring within the same biogeographic region. Here, we use molecular data in the form of ultra-conserved elements (UCEs) to infer the evolutionary relationships and population genomic structure of the Eastern Pinesnake complex (*Pituophis melanoleucus*) – a polytypic wide-ranging species that occupies much of the Eastern Nearctic. In addition to inferring evolutionary relationships, population genomic structure, and gene flow, we also test relationships between genomic diversity and putative barriers to dispersal, environmental variation, and geographic distance. We present results that reveal shallow population genomic structure and ongoing gene flow, despite an extensive geographic range that transcends geographic features found to reduce gene flow among many taxa, including other squamate reptiles within the Eastern Nearctic. Further, our results indicate that the observed genomic diversity is spatially distributed as a pattern of isolation by distance and suggest that the current subspecific taxonomy do not adhere to independent lineages, but rather, show a significant amount of admixture across the entire *P. melanoleucus* range.

## Introduction

Understanding the processes and underlying mechanisms that promote and maintain biological diversity (i.e., speciation) is a central goal in evolutionary biology (De Queiroz, 1998; Turelli, Barton, & Coyne, 2001; Wiley & Lieberman, 2011). For example, the most widely acknowledged mode of speciation is one in which populations diverge due to geographic isolation (allopatric speciation), allowing for the accumulation of fixed genetic differences over time (Hoskin, Higgie, McDonald, & Moritz, 2005; Mayr, 1963). Most often, geographic barriers to dispersal and environmental variation (i.e., isolation by environment [IBE]) underpin these reductions in gene flow, and are therefore important to identify when attempting to understand the evolutionary histories of diverged lineages. However, genomic variation may be distributed across geographic space as clines (i.e., isolation by distance [IBD]) when barriers to dispersal or environmental variation only minimally reduce gene flow (Slatkin, 1993; Wright, 1943). In instances where species have greater dispersal capabilities and encounter only weak barriers to gene flow, genomic variation can also be uniformly distributed across panmictic populations; this being most frequently observed in marine and/or migratory taxa (Coltman et al., 2007; Palm, Dannewitz, Prestegaard, & Wickström, 2009; White, Fotherby, Stephens, & Hoelzel, 2011). Although examples of divergence with gene flow are becoming increasingly common (Feder, Egan, & Nosil, 2012), it may still be more likely that gene flow among demes would homogenize genetic differences and ultimately prevent lineage divergence (Bolnick & Fitzpatrick, 2007; Dobzhansky & Dobzhansky, 1971; Endler, 1973; Mayr, 1963). As such, it may be equally valuable to identify when genetic continuity has been maintained across a species’ geographic distribution, especially when a species occupies an ecologically complex and expansive region with previously recognized biogeographic barriers.

Species with distributions that span large biogeographic regions are more likely to encounter barriers to gene flow and environmental variation that can lead to population genomic structuring compared to species inhabiting more homogenous landscapes (Avise et al., 1987; Soltis, Morris, McLachlan, Manos, & Soltis, 2006). For example, the Eastern Nearctic is a large biogeographic region that spans much of North America and harbors multiple biodiversity hotspots and several well-known biogeographic barriers that reduce gene flow among populations of different species in predictable ways (Burbrink & Guiher, 2015; Burbrink, Lawson, & Slowinski, 2000; McKelvy & Burbrink, 2017; Myers, McKelvy, & Burbrink, 2020). As such, previous studies have shown that the population genomic structure of squamate reptiles within this region tends to correlate with the Apalachicola and Mississippi river drainages (Alexander Pyron & Burbrink, 2009; Burbrink et al., 2000; Weinell & Austin, 2017). However, the impacts of biogeographic heterogeneity on population genetic structure may be taxon specific (as reviewed by Soltis et al., 2006), depending on species range distributions, genetic diversity, and overall vagility. Species with considerable dispersal abilities, such as large mammals (DeCesare et al., 2020; Latch, Heffelfinger, Fike, & Rhodes Jr, 2009) and some birds (Kekkonen et al., 2011), might be expected to exhibit more homogenized levels of genetic diversity than species with more limited vagility. Even within squamate reptiles, different mechanisms may underpin the population structure of species with different dispersal capabilities. Investigating links between landscape features and genomic diversity of widespread taxa within the Eastern Nearctic or similar regions may therefore refine our understanding of the processes and underlying mechanisms that promote and maintain biological diversity.

The Eastern Pinesnake complex (*Pituophis melanoleucus*) is a polytypic wide-ranging species complex occupying much of eastern North America with several disjunct populations. This species typically inhabits upland pine ecosystems, has been categorized as a longleaf specialist in the most southern regions of its distribution (Baxley & Qualls, 2009), and has one of the largest home range sizes known across colubroid snakes (Miller, Smith, Johnson, & Franz, 2012). Color patterns vary across its range; from being uniformly black in areas of southern Mississippi and Alabama, to a combination of white and dark blotches of red and bronze in the south/central distribution and black and white in the New Jersey Pine Barrens (Guyer, Bailey, & Mount, 2019). The current taxonomic classification of this complex – largely based on color pattern, recognizes three geographical sub-specific taxa: (1) the Northern Pinesnake (*P. m. melanoleucus*), (2) the Florida Pinesnake (*P. m. mugitus*), and (3) the Black Pinesnake (*P. m. lodingi*) (Crother, 2012). All three subspecies are of taxonomic interest because of conservation concerns stemming from habitat degradation/fragmentation and declining prey communities (e.g., southern pocket gopher decline, longleaf pine degradation; (Landers, Van Lear, & Boyer, 1995; Van Lear, Carroll, Kapeluck, & Johnson, 2005). Previous molecular studies that included members within this complex have not incorporated population scale sampling and resulted in variable hypotheses regarding placement within the genus *Pituophis* (Bryson Jr, García‐Vázquez, & Riddle, 2011; Pyron & Burbrink, 2009; Rodrı́guez-Robles & De Jesús-Escobar, 2000). Because of the current ambiguous relationships, lack of geographic structuring recovered from traditional molecular data, morphological variability, conservation concern, and vast geographic distribution, this group represents an ideal model to investigate phylogenetic and population genomic patterns. Moreover, this species complex provides additional opportunity to continue testing hypotheses regarding population differentiation and known geographic barriers. Here, we aim to quantify population genomic structure within the *P. melanoleucus* complex and determine if spatial patterns of genomic variation result from barriers to gene flow, environmental variation, geographic distance, or continuous gene flow across the entire geographic range. To this end, we use nuclear genomic data sampled across the entire geographic range in the form of ultraconserved elements (UCEs) to (1) infer the evolutionary relationships and population structure within the *P. melanoleucus* complex, (2) test for barriers to gene flow, IBE, and IBD, and (3) address the taxonomic validity of the currently recognized sub-specific designations. Notably, this is the first population genomic study of this species, providing invaluable information regarding levels of genetic differentiation and insight for future conservation genomic efforts.

## Methods and Materials

### Sample Collection and DNA Extraction

Forty-three tissue samples were collected from individual specimens of *Pituophis melanoleucus*, including all of the recognized subspecies across its geographic distribution with five additional samples collected from specimens of *P. ruthveni, P. catenifer,* and *Pantherophis obsoletus* to serve as outgroup taxa. Tissue samples were taken either from liver, muscle, shed skins or ventral scale clips and preserved in 95-100% ethanol until DNA extraction. Genomic DNA was isolated by using Qiagen’s DNeasy kit following the manufacturers protocols and quantified using a Qubit 2.0 (Thermofisher Scientific, Waltham, Massachusetts, USA). DNA samples were sent to the University of Georgia’s Department of Genetics for library preparation and sequencing of (UCEs).

### UCE assembly and dataset generation

Samples were barcoded using Illumina TruSeq adapters with unique 8bp sequence tags for each individual and UCEs were targeted by using a tetrapod 5060-locus probe set (open source availability from https://ultraconserved.org). Demultiplexed samples were filtered and processed by removing adapter sequences and ambiguous bases using the program Illumiprocessor which is incorporated in the open source software package Phyluce v.1.5 (Faircloth, 2016). Reads were assembled *de novo* using standard settings in Velvet v.1.1 (Zerbino & Birney, 2008) and assembled contigs were matched against the 5k UCE tetrapod probe kit to identify and extract UCEs within Phyluce. Sequence alignments for each individual UCE locus were generated using MAFFT v.7.397 (Katoh & Standley, 2013) and concatenated data alignment matrices of 50% and 75% were created in order to examine the effects of missing data on resulting topologies. The data matrix percentages represent the amount of alignments that include every individual (e.g., 50% data matrix would indicate that all alignments will contain at least half of the individuals). Python scripts within Phyluce were used to generate summary statistics of alignment matrices, total reads, contigs, and UCE counts.

To call variants from our assembled sequence capture data, we used the pipeline outlined in Harvey, Smith, Glenn, Faircloth, & Brumfield, 2016, which uses samtools (Li et al., 2009) and Picard (http://picard.sourceforge.net) to convert, merge, and index BAM files and remove PCR duplicates. We then used the best practices workflow from GATK (McKenna et al., 2010) to call and hard filter biallelic SNPs that had low quality scores and read depth (i.e., GQ<30 and DP<5). To avoid effects of linkage and violating underlying assumptions for the population genetic clustering software used, we sampled only the first SNP from each UCE loci.

### Phylogenetic Analysis and Population genomic analyses

We inferred phylogenetic hypotheses in a maximum-likelihood (ML) framework using the program RAxML v.8.2.3 (Stamatakis, 2014). Twenty ML searches were performed on each dataset (50%, 75%, and first SNP dataset) by partitioning each individual UCE locus under the GTR+Γ model with the exception of the SNP dataset. This partitioning scheme was chosen over a single partition to estimate individual locus specific parameters (e.g., base frequencies, transition/transversion ratios). Node support was assessed by performing 100 BS replicates for each dataset and visualized using the program FigTree v1.4.4 (http://tree.bio.ed.ac.uk/software/figtree/).

To understand and infer overall population structure and genetic clustering we first performed a principal component analyses (PCA) in the R package adegenet (Jombart & Ahmed, 2011) to visualize the spatial distribution of genetic variance. We then used the program STRUCTURE v.2.3.4 (Falush, Stephens, & Pritchard, 2003; Hubisz, Falush, Stephens, & Pritchard, 2009; Pritchard, Stephens, & Donnelly, 2000) to infer levels of gene flow using the admixture model and correlated allele frequencies with no prior assumption of individual or population assignments. The program Strauto (Chhatre & Emerson, 2017) was used to run multiple iterations of *K* values (2-7) by utilizing the parallelization function. For each *K* value, 20 iterations of 10,000 burn-in generations were conducted followed by 50,000 logged generations. To determine the optimal value of *K,* the Δ*K* (Evanno, Regnaut, & Goudet, 2005) values were evaluated by using the program STRUCTURE HARVESTER (Earl, 2012). The program CLUMPP was used to combine all iterations for each *K* value using the greedy algorithm to estimate the most likely membership coefficients. The results were visualized using the package pophelper in R (Francis, 2017).

In addition to STRUCTURE, we also used R package conStruct (Bradburd et al., 2018) to assess population genomic structure with regard to geographic distance. The program conStruct simultaneously infers discrete (non-spatial model) and continuous (spatial model) patterns of population genomic structure by estimating ancestry coefficients for each sampled individual from two-dimensional population layers. Using a cross-validation procedure, conStruct is then able to assess differential support for both non-spatial and spatial models. Spatial models are given greater support by conStruct when the decay of allelic covariance is strongly associated with increasing geographic distance. In such cases, conStruct can then assign all individuals to a single population layer, where other clustering algorithms cannot, overestimating the number of discrete clusters (Bradburd et al., 2018; Frantz, Cellina, Krier, Schley, & Burke, 2009; Meirmans, 2012). Given that conStruct does not perform well with sparse data matrices (Bradburd et al., 2018), we further filtered our original SNP dataset using VCFtools (Danecek et al., 2011), to include only sites that were present in at least 75% of samples and excluded 3 samples that had high degrees of missing data, resulting in 1876 SNPs. We then used the cross-validation method to determine the best fit model between both spatial and non-spatial models for *K* values of two through six with ten replicates per *K* and 25,000 iterations per replicate with a training proportion of 0.75. We determined the optimal value of *K* by comparing the resulting predictive accuracy of each model with the individual layer contributions, testing for differences with a standard t-test in R base package. We then ran five independent conStruct analyses for both non-spatial and spatial models at each *K* value with 25,000 iterations per and confirmed convergence of analyses by analyzing trace plots in conStruct. Last, we tested correlations between ancestry coefficients estimated between STRUCTURE and a non-spatial model in conStruct for *K*=2 using a Pearson’s correlation. Outputs from both STRUCTURE and construct were then plotted in R using a combination of the conStruct package and custom scripts (Bradburd et al., 2018).

### Estimation of Effective Migration Surface

To visualize patterns of gene flow across the *P. melanoleucus* range, we used the program EEMS – Estimation of Effective Migration Surface (Petkova, Novembre, & Stephens, 2016), which estimates effective migration and provides visualization of migration surfaces that highlight geographic regions of relatively increased and reduced gene flow. EEMS uses a stepping-stone model to relate effective migration rates to expected genetic dissimilarities of geo-referenced samples. We estimated gene flow using three independent chains, each with 750 demes, for 1,000,000 MCMC iterations, with 50,000 iterations of burn-in and a thinning interval of 1,000. We plotted results to check for agreement across all three independent chains and checked for MCMC convergence by visualizing trace plots using the R package, rEEMSplots (Petkova et al., 2016).

### Generalized Dissimilarity Modeling

In order to test the roles of biogeographic barriers, environmental variation, and geographic distance in shaping genomic variation across the *P. melanoleucus* range, we used generalized dissimilarity modeling (GDM) on a reduced set of BioClim variables, elevation, geographic distance, and several river drainages using R package gdm (Manion et al., 2017). In particular, we included the Alabama river system, the Apalachicola river, and the Suwannee river, as these river systems have been previously shown to reduce gene flow between demes of several small vertebrate taxa (Avise, Giblin-Davidson, Laerm, Patton, & Lansman, 1979; Burbrink et al., 2000; Church, Kraus, Mitchell, Church, & Taylor, 2003; Jackson & Austin, 2010; Walker & Avise, 1998). Generalized dissimilarity modeling is a permuted matrix regression technique that has gained recent popularity in population genomic studies for parsing relationships between geographic features, geographic distances, environmental distances, and genetic distances. Briefly, GDM uses a permuted nonlinear matrix regression through the use of I-spline basis functions to accommodate nonlinear statistical relationships (Manion et al., 2017) and incorporates backwards model selection to assess relationships between the response and individual predictor variables.

We downloaded the complete set of BioClim variables at a resolution of 2.5 minutes using the R raster package and removed variables with correlations of 0.75 or greater with the *raster.cor.matrix* function in ENMtools R package (Warren, Glor, & Turelli, 2010), as in Myers et al. (2019). Elevation for each sampling point was also acquired with the raster package using the function *get_elev_points*. Variable reduction retained seven BioClim variables (mean diurnal temperature range, maximum temperature of the warmest month, mean temperature of the wettest quarter, mean temperature of the driest quarter, annual precipitation, precipitation of the driest month, and precipitation o the warmest quarter), which were set as predictor variables in our models. River drainages were determined using custom R scripts which assigned samples to different “basins” according to their orientation to different river systems. For example, samples located west of the Alabama river were assigned a value of four while samples west of the Suwanee River but east of the Apalachicola River were assigned a value of two. We then calculated the ‘distances’ between samples where a value of one would represent samples being separated by a single river system, a value of two represented samples separated by two river systems, and so forth. Our response variable, Euclidean genetic distance, was estimated from our original VCF using the *dist* function in R package adegenet (Jombart & Ahmed, 2011), using the ‘Euclidean’ flag.

We used GDM to assess the relationships between all predictor variables and the response variable with three I-spline knots and 100 jackknife replicates as in Fitzpatrick & Keller (2015). For each replicate, we removed all samples from a random 10% of sampling locations and model fit was assessed according to the percent deviance explained between permuted and un-permuted models. In order to assess the importance of each individual variable (each of the seven BioClim variables, geographic distance, elevation, and the river basins), we used backwards elimination by running the *gdm.varimp* function in the gdm package (Ferrier, Manion, Elith, & Richardson, 2007; Manion et al., 2017) with 100 permutations for each step. This function determines significance for each predictor variable by permuting predictor variables individually and assessing the deviance explained between the un-permuted model and the different permuted models. The relationships between genetic distance and the predictor variables that were determined significant through backwards elimination of the full model were then plotted with the R package gdm (Ferrier et al., 2007) using the *plotUncertainty* function after running a second, refined model using only these significant predictor variables. Moreover, we once again ran the *gdm.varimp* function in the gdm packaged to assess the relative contribution of each of the predictor variables included in our refined model (geographic distance, mean diurnal range, precipitation of the warmest quarter, and river basin).

## Results

### Sequence data and data matrices

The average number of reads for each individual was 1,886,033 with a contig range of 7,136 to 19,784 (Table S1). The number of recovered UCEs from all individuals ranged from 3,383 to 4,156 with an average length of 596 bp. The 50% data matrix contained 4,332 UCEs with a total of 2,768,605 bp and the 75% alignment had 3,996 UCEs and 2,623,726 bp. From our variant calling pipeline we called a total of 2910 SNPs that were used in all subsequent population clustering analyses with the exception of conStruct in which we filtered down to include 1876 SNPs. For phylogenetic analyses using only SNP data, which included outgroup taxa, we retained 2715 SNPs that passed all hard filtering parameters.

### Phylogenomic analyses

Our phylogenetic results across all datasets (50%, 75%, and SNP) recovered *P. melanoleucus* as a monophyletic group and sister to *P. ruthveni* (Bootstrap value of 100; Figure 1a and Supplemental Figure 1). All inferred topologies did not recover the currently recognized subspecific taxa as monophyletic and generally held the same geographic pattern with samples from the extreme edges of the species distribution tend to group together with high node support (Bootstrap value > 70; e.g., North Carolina/New Jersey, Mississippi, and Peninsula Florida; Figure 1a). Samples that were located in the central region showed the greatest shift in subclade membership as the topology placement was the most variable for locations in central Alabama, Georgia, and panhandle of Florida (e.g., Covington, AL, Marion, GA, and Hernanodo, FL; Supplemental Figure 1). Node support varied across topologies with most receiving low support values (<70), with the exception of the subclade of all North Carolina and New Jersey individuals and the two Tennessee individuals within the 50 and 75% topologies.

**Figure 1.**
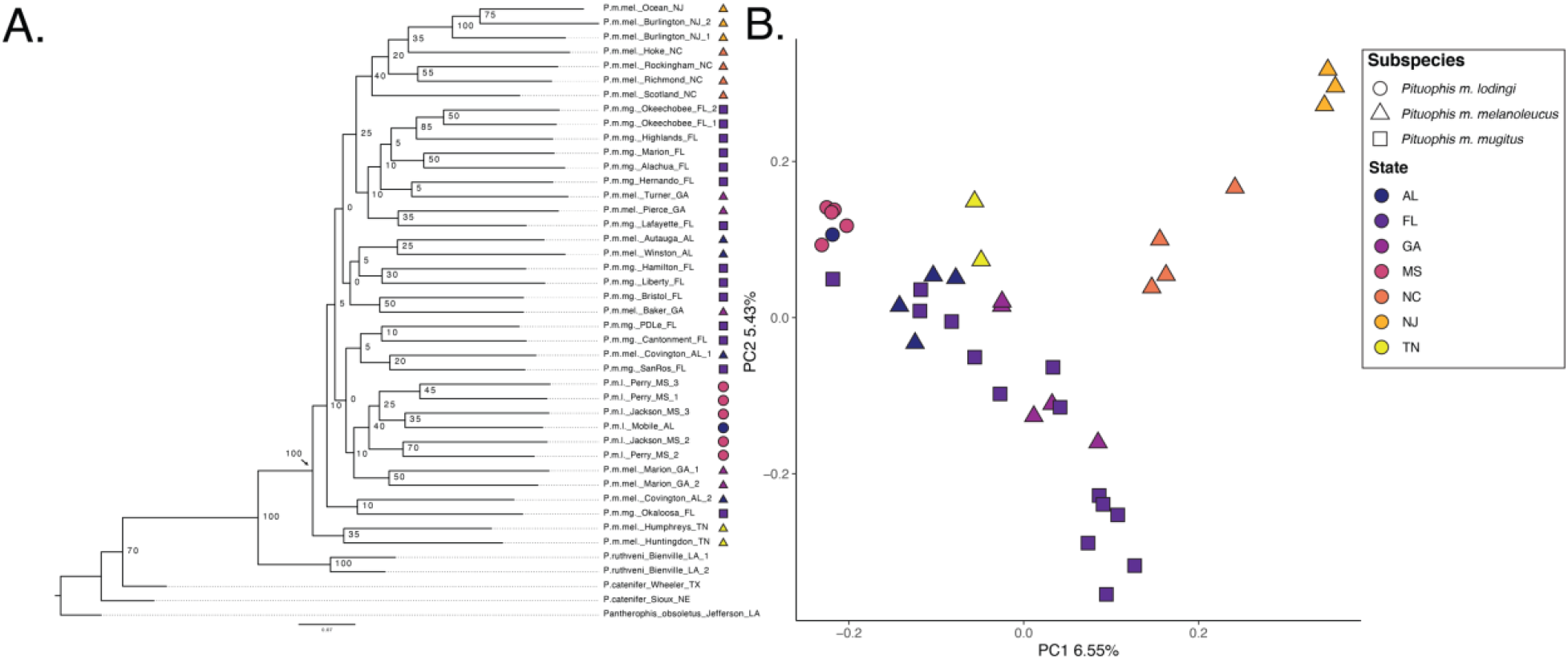
Panel highlighting the genetic structuring and patterns across the Eastern Pinesnake complex. A. Results of the maximum likelihood SNP tree with tips and shapes corresponding to the subspecific taxonomic classification and state locality which is also highlighted in the subsequent panel. B. Principle coordinate analysis (PCA) plot showing the distribution of genetic variance of all individuals for the first two loadings.

### Population Cluster analysis

The final dataset used in all population clustering analyses (with the exception of the conStruct analysis) contained a total of 2,910 SNPs with the amount of individual missing data ranging from 1-58% (average:13%). Principal component analysis revealed a pattern of continuous variation with PC1 describing 6.55% of the variation in the data, and PC2 describing 5.43%. Notably, PCA revealed overlap between all three recognized subspecies classifications (Figure 1b). Results from our STRUCTURE runs showed that the best supported model using the Δ*K* method was a *K* value of 2 (Supplemental Figure 2) with ancestry coefficients/admixture levels exhibiting a gradient distributed from east to west across the Florida panhandle and Apalachicola River drainage (Figure 2a). This clinal gradient was further distributed from North Florida through the Piedmont and into central Tennessee. Samples along the East coast in North Carolina and New Jersey were maintained within the cluster assigned to samples throughout the Florida peninsula. Shifts between majority proportions of ancestry occurred at the Alabama river drainage, discretely clustering samples east and west of this drainage. Like STRUCTURE, conStruct also revealed a gradient of ancestry coefficients and gave greatest support to a model with two population layers (t = 10.46; p < 0.001) (Figure 2b). Notably, for *K* = 2, ancestry coefficients estimated with a non-spatial model in conStruct were highly correlated with the ancestry coefficients estimated by STRUCTURE (correlation = 0.98; p < 0.001).

**Figure 2.**
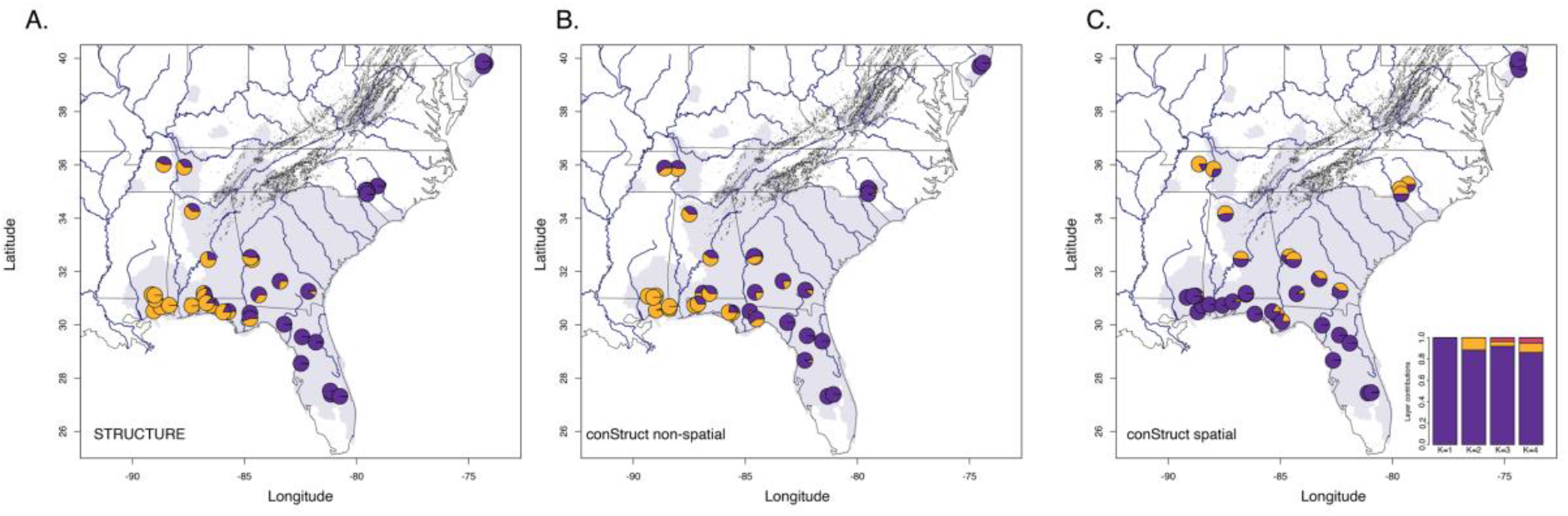
Geographic patterns of ancestry coefficients. A., B., and C. Map of samples with individuals shown as pie charts for *K*=2 model. Colors represent cluster membership across STRUCTURE (A) and conStruct (both non-spatial and spatial models; B. & C) analyses, which are indicated in the bottom left corner of each panel. Panel C inset shows the dissection of layer contributions for conStruct spatial models sampled across *K* values ranging from one to four.

Observations of clinal genomic variation were further supported by the cross validation of non-spatial and spatial models in conStruct, which gave greater support for spatial models than for non-spatial models at all values of *K* (Supplemental Figure 3). The preference of a spatial model indicates an underlying pattern of isolation by distance (IBD), where allelic covariation is associated with geographic distance. The analysis of layer contributions then suggested that little resolution in the genomic data was described by adding even a second layer (Figure 2c, inset). This inference is apparent in our visualization of ancestry coefficients generated through a spatial model in conStruct plotted atop a geographic map, which showed a majority population layer that persisted across the entire *P. melanoleucus* range (Figure 2c). Unlike STRUCTURE and non-spatial conStruct models, visualization of ancestry coefficients generated by the spatial model of *K* = 2 by conStruct showed very little population genomic structure occurring around any river system. Indeed, the greatest population genomic structure according to this model appears to have occurred throughout the core of the *P. melanoleucus* range, which roughly corresponds to the Piedmont region.

Last, a conStruct non-spatial model of *K* =3 showed population genomic structuring that occurred at the Alabama river drainage, along with a third population layer that became evident at higher latitudes (e.g., Tennessee, North Carolina, and New Jersey) (Supplemental figure 4a). However, adding a third population layer in the spatial conStruct model did not show any population genomic structuring that occurred around any river system, and only began to add this third layer into samples from the Piedmont region (Supplemental figure 4b).

### Estimation of Effective Migration Surface

Visualization of Estimation of Effective Migration Surface (EEMS) is an additional method used to assess gene flow and population genomic structure with respect to geographic features. Our results from EEMS (Figure 3a) revealed patterns of gene flow that were consistent with other population genomic analyses presented here (Figures 1 and 2), where no clear barrier to effective migration was highlighted. Indeed, there appears to be relatively consistent effective migration across the entire *P. melanoleucus* range, which would reduce any population genomic stratification. Notably, the only region of reduced effective migration highlighted by EEMS occurs throughout much of the Piedmont, where conStruct models indicate increased population genomic structure.

**Figure 3.**
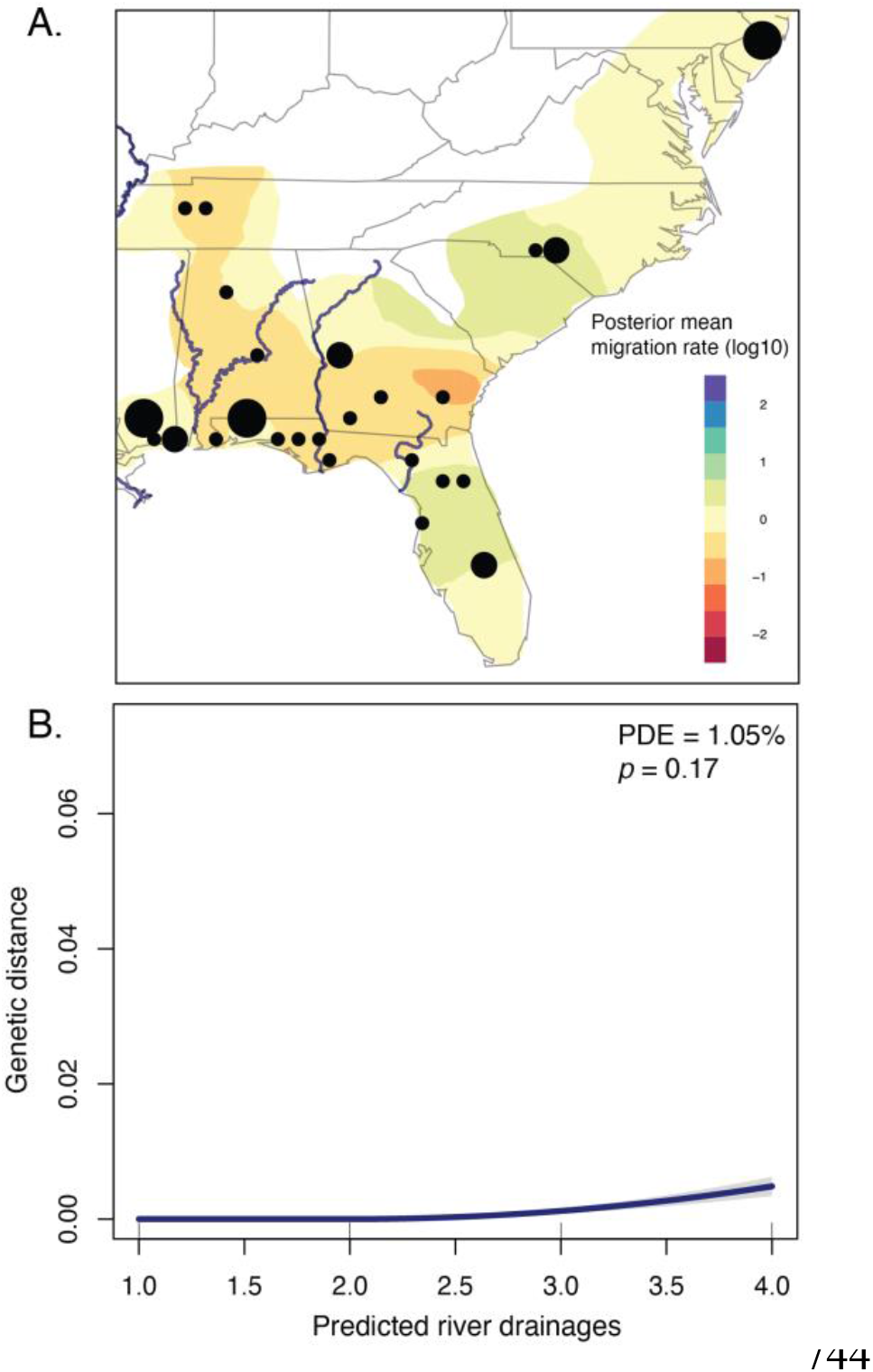
Estimates of effective migration surfaces (EEMS) and correlations of genetic distance across major river basins. A. EEMS plot showing the migration surface for all sampled individuals. Reduced rates of effective migration are represented by red and yellow shades while blue and purple shades reflect increased rates of effective migration. B. Output from generalized dissimilarity modeling showing the relationship between pairwise genetic (y-axis) distances and the predicted number of river basins separating samples (x-axis). Note that the percent deviance explained and the p-value for the model are in the top right corner and show the variation in genetic distance explained by the predictor variable does not significantly deviate between unpermuted and permuted models.

### Generalized Dissimilarity Modeling

Last, to test for biogeographic barriers to gene flow, as well as patterns of isolation by environment (IBE), and IBD, we used generalized dissimilarity modeling (GDM). Results of GDM indicated (Table S2: Full model and Table 2: refined model) that the percent deviance explained by the full model that included all seven BioClim variables, geographic distance, elevation, and the river basins was 46.7% (p < 0.001). However, a refined model incorporating only the predictor variables determined to be significant in the full model (geographic distance, mean diurnal temperature range, precipitation of the warmest quarter, and river basins) still accounted for 46.3% (p < 0.001) of the deviance explained.

Backwards elimination of predictor variables within the refined model further indicated that the river basins only account for 1.1% of the deviance explained from the refined model (Figure 3b) and were ultimately not significant predictors of variation in genetic distance (p = 0.17). Mean diurnal range and precipitation of the warmest quarter, meanwhile, explained 22.6% (Figures 4a and 4b) (p = 0.03) and 11.9% (Figures 4c and 4d) (p = 0.02) deviance from the full model, respectively. However, geographic distance accounted for 63.2% (Figure 4e) of the deviance explained from the full model (p < 0.001), lending statistical support for a model of IBD over geographic barriers, IBE, or panmixia.

**Figure 4.**
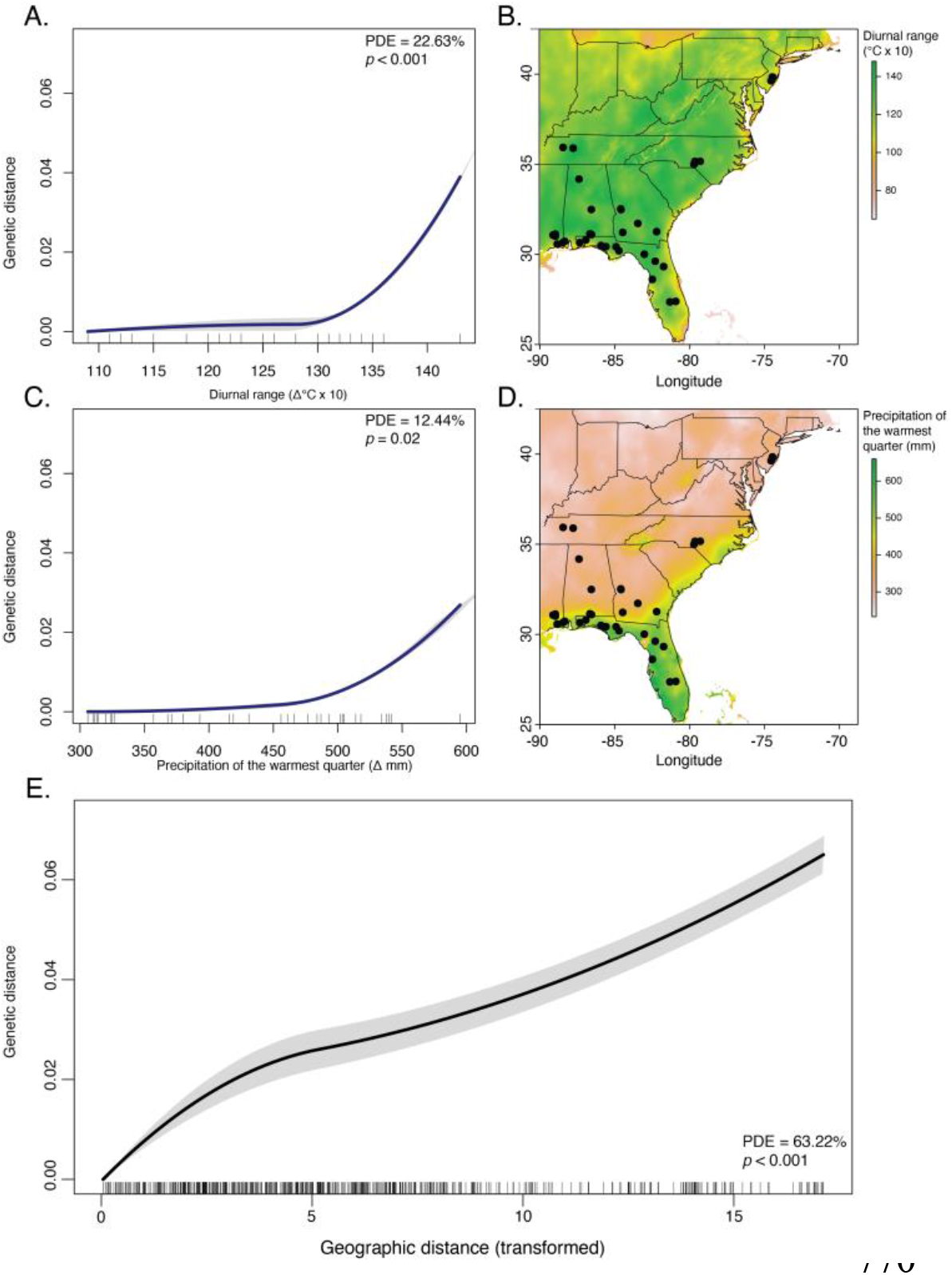
Results from generalized dissimilarity modeling and maps of environmental variation across the *Pituophis melanoelucus* range. A. The relationship between pairwise genetic distance and variation in mean diurnal monthly temperature range (degrees Celsius x 10 [max – min]) estimated from generalized dissimilarly modeling. B. Mean diurnal monthly temperature range (degrees Celsius x 10 [max – min]) across the *P. melanoleucus* range. C. The relationship between pairwise genetic distance and variation in precipitation of the warmest quarter (mm) from generalized dissimilarly modeling. D. Precipitation of the warmest quarter (mm) across the *P. melanoleucus* range. E. The relationship between pairwise genetic distance and transformed geographic distance estimated with generalized dissimilarity modeling. The percent deviance explained by the permuted model from the unpermuted model and the associated p-value are in the top right or bottom right of each plot from our generalized dissimilarity modeling output.

## Discussion

### Evidence for a single lineage

Here, we present the first phylogenetic and population genomic-scale study of the Eastern Pinesnake (*Pituophis melanoleucus*) species complex. Our results indicate that the currently recognized subspecific nomenclature does not reflect evolutionary history as subspecies were not recovered as independent lineages (i.e., monophyletic) (Figure 1 and Supplemental Figure 1). Rather, this complex appears to be a single species composed of multiple subpopulations with little genetic differentiation indicated by shallow population genomic structuring and moderate levels of gene flow across the entire species distribution, which contrasts many of the previously studied squamate taxa within the Eastern Nearctic (Soltis et al., 2006).

Results from our principal components analysis (PCA) revealed a pattern of genetic diversity that anecdotally resembled our geographic sampling distribution and showed no obvious clustering between sub-specific taxonomic designations (Figure 1b). Such patterns in PCA (and lack of monophyly) tend to reflect genetic diversity that is continuously distributed across species’ ranges with no phylogenetic breaks and little population stratification (Novembre & Stephens, 2008; Reich, Price, & Patterson, 2008). Results from our STRUCTURE and conStruct analyses also show a clinal pattern of genetic diversity where ancestry coefficients appear to be continuously distributed across geographic space (Figures 2a-c). Moreover, cross validation in conStruct supports a spatial model over a non-spatial model at all values of *K*, indicating a strong correlation between genetic and geographic distance. Combined with results from our tree-based approach, our clustering analyses suggest a pattern of genetic variation would be consistent with IBE or IBD (Wang & Bradburd, 2014), and not biogeographic barriers or panmixia, which would likely lead to either monophyly and multiple discrete clusters or a single population layer. Although STRUCTURE (Figure 2a) and non-spatial conStruct models (Figures 2b and Supplemental Figure 4a), which are highly correlated, reveal population genomic structure (*K*=2) that occurs near the Western edge of the *P. melanoleucus* range at the Alabama river drainage, this is likely an artefact of the observed clinal variation and not evidence of a biogeographic barrier. Spatial conStruct models (which are supported over non-spatial models) do not indicate population stratification that occurs near any river system, but rather show shallow structure that occurs throughout the central region of the *P. melanoleucus* range (i.e., the Piedmont) (Figure 2c and Supplemental Figure 4b), which is likely due to genetic drift (Bradburd et al., 2018), potentially brought on by habitat fragmentation (Maloney & Weller, 2011; Taverna, Peet, & Phillips, 2005). Indeed, differences between non-spatial and spatial clustering models presented here may even begin to highlight the importance of incorporating a spatial term in clustering algorithms, as phylogenomic breaks can be prematurely inferred from genomic gradients that transcend potential biogeographic barriers. Here, visualization of ancestry coefficients estimated with spatial conStruct models reveals a majority layer contribution that occurs across the entire *P. melanoleucus* range. Layer contribution analysis implemented in conStruct also shows that little additional resolution in the genomic data is gained by adding layers above *K* =1 (Figure 2c inset). Collectively, these results suggest that geographic barriers have had little to no impact on historical population genomic structure within this species complex and that gene flow across the *P. melanoleucus* range and current subspecific designations is likely ongoing.

### Gene flow across a previously recognized genetic barrier

Like clustering analyses, visualization of effective migration across the *P. melanoleucus* range using EEMS shows a moderate reduction in gene flow within Piedmont region, but not adjacent to any river systems (Figure 3a). Rather, effective migration across the entire *P. melanoleucus* range appears to be moderate, with even slightly increased effective migration in parts of South Carolina and Florida. These results contrast previous squamate phylogeography studies (Burbrink, Fontanella, Pyron, Guiher, & Jimenez, 2008; Burbrink et al., 2000; Weinell & Austin, 2017, Myers et al., 2020) which tend to show substantial genetic variation between populations and clade memberships occurring on opposite sides of different river drainages. Discord among results from our phylogenetic and population genomic structuring analyses compared to other squamate taxa within the Eastern Nearctic may feasibly be underpinned by the dispersal capabilities of these species. Previous studies of movement patterns and annual home range sizes within the genus *Pituophis* span from approximately 35 hectares in *P. catenifer* (Kapfer, Pekar, Reineke, Coggins, & Hay, 2010), to 70 hectares in *P. m. mugitus* (Miller et al., 2012), and over 105 hectares in *P. m. melanoleucus* (Zappalorti, Burger, & Peterson, 2015). Conversely, home range estimates for *Pantherophis* tend to converge on a mean of approximately 10 hectares in *P. alleghaniensis* (Carfagno & Weatherhead, 2008; Durner & Gates, 1993) and 50 hectares in *P. gloydi* (Row, Blouin-Demers, & Lougheed, 2012). Additionally, annual range estimates within *Lampropeltis* have been documented at approximately 15 hectares for *L. getula* (Wund, Torocco, Zappalorti, & Reinert, 2007) and eight hectares for *L. calligaster* (Richardson, Weatherhead, & Brawn, 2006). Because dispersal capability is often directly proportional to home range size (Choudoir, Barberán, Menninger, Dunn, & Fierer, 2018; Pérez‐Pérez, López‐Moreno, Suárez‐Rodríguez, Rheubert, & Hernández‐Gallegos, 2017) and inversely related to genetic structure (Brunke et al., 2020; Epperson & Alvarez‐Buylla, 1997; Polato et al., 2018; Wright, 1943), it is possible that the larger home range size estimates in *P. melanoleucus* accounts for the lack of discreet phylogenetic breaks that are commonly observed in snake species within the Eastern Nearctic.

### Isolation by Environment and Isolation by Distance

In addition to barriers to gene flow, environmental variation can also structure genomic diversity between populations and lead to discreet phylogenetic breaks (Coyne & Orr, 2004). However, environmental variation can also shape genetic diversity across species’ ranges as clines, which may be difficult to distinguish from patterns of IBD (Lee and Mitchell-Olds, 2011). Using generalized dissimilarity modeling (GDM), which holds individual predictor variables constant, we simultaneously tested hypotheses of barriers to gene flow, IBE, IBD, and panmixia. Results from our GDM analysis support the hypothesis that continuous gene flow has not been impacted from river drainages within the *P. melanoleucus* complex as these features explain only 1.1% of the deviance and it is not significant (Figure 3b). Using backwards elimination implemented in GDM, we also found moderate but significant patterns of IBE (Figures 4a-4d). Additionally, we find that the range of diurnal temperatures across the year, as well as the amount of precipitation during the warmest quarter of the year, describe 22.6 and 11.9% of the deviance from our refined model with the associations between genetic distance and environmental variation to be the greatest when environmental differences were high. This may explain the variation in ancestry coefficients within our genomic dataset, such as those observed at the transition between the Piedmont and Coastal Plains regions (Figures 2a-2c and Supplemental Figures 4a and 4b).

Although this environmental variation could be linked to local adaptation, potentially even for morphological traits such as color pattern (Rosenblum, 2006), it may also drive variation in movement patterns and thus, rates of gene flow between the demes of different regions. However, we still found greatest support for a pattern of IBD within *P. melanoleucus*, where geographic distance described over 62% of the variation in our genomic data (deviance from the refined model) (Figure 4e). This correlation indicates that *P. melanoleucus* is not a truly panmictic species, which would be represented by a weak association between genetic and geographic distances. These results are also concordant with previous findings that ectothermic species are less likely to be panmictic than other terrestrial vertebrates due to metabolic and energetic requirements (Jenkins et al., 2010). However, among the squamate reptiles of the Eastern Nearctic, *P. melanoleucus* appears to be unique in that river systems have had little to do with shaping genomic diversity across its range.

## Conclusion

Our analysis of genomic diversity, population genomic structuring, and gene flow provides insight into how widespread, phenotypically distinct species complexes may have traditionally been regarded as separate lineages, yet these complexes exhibit significant levels of genetic homogenization. More specifically, our results highlight that geographic features typically considered to be biogeographic barriers, can have varying impacts on the population genomic structure of different taxa. It may therefore be naïve to accept certain taxa as paradigms for phylogeographic studies, even among closely related species. Here, genomic variation across the *P. melanoleucus* range is best describe by IBD, as individuals of proximate demes are more likely to exchange alleles than individuals of more geographically distant localities. It is this relationship between geographic distance and genetic decay that has ultimately shaped the shallow and continuous population genomic structure observed across the *P. melanoleucus* range. Although genomic patterns of IBD can lead to reproductively-isolated lineages through either stochastic “breaks” (Baptestini, de Aguiar, & Bar-Yam, 2013; Hoelzer, Drewes, Meier, & Doursat, 2008) or “speciation through forced distance” (i.e., ring species; Irwin, & Price, 2005), this is unlikely to be the case here provided the lack of reciprocal monophyly and shallow levels of population genomic stratification.

While future studies focusing on intraspecies level relationships within the genus *Pituophis* would benefit from increased sampling, the results observed here provide additional evidence for the current lack of understanding regarding the evolutionary relationships among these snakes and suggests that currently this species complex appears to be comprised of a single lineage. We acknowledge that molecular markers other than UCEs may be better suited for these analyses, however, given the inferred shallow nature of this species complex, UCEs are likely still viable for our data and even benefit because they do not suffer from lineage specific dropout as opposed to other reduced representation methods.

## Acknowledgments

We would like to thank the following institutions and people for providing samples for this project: the Florida Museum of Natural History (K. Krysko, and K. Enge) Louisiana State University (D. Dittman), North Carolina Museum of Natural Sciences (J. Beane), the Nature Conservancy (J. Lee), American Museum of Natural History (F. Burbrink and A. Mckelvy), Auburn University (B. Folt and D. Laurencio), and the University of Southern Mississippi (B. Kreiser). This research was funded by the Schlieder Foundation.

## Data accessibility

Raw data has been deposited under the BioProject number: PRJNAxxxxxx

---

**Table 1.** Results from generalized dissimilarity modeling (gdm) including backwards elimination of individual predictor variables for the refined model. The full refined model includes the four predictor variables determined to be significant in the original full model (Table 1). Each successive model includes all variables from the previous model barring the predictor variable with the least contribution to percent deviance explained. Significance was determined in gdm through 100 permutations. Note that we highlighted our final accepted model that included a model where all predictor variables were determined to be significant. This final model included geographic distance, precipitation of the warmest quarter, and mean diurnal temperature range.

| Predictor variable | Full model |  | Model two |  | Model three |  |
| --- | --- | --- | --- | --- | --- | --- |
|  | % deviance | P value | % deviance | P value | % deviance | P value |
| Geographic distance | 24.31 | 0.00 | 63.22 | 0.00 | 63.25 | 0.00 |
| Mean diurnal temperature range | 21.40 | 0.00 | 22.63 | 0.00 | 22.70 | 0.03 |
| Precipitation of the warmest quarter | 12.44 | 0.01 | 11.93 | 0.02 | NA | NA |
| River basin | 1.05 | 0.17 | NA | NA | NA | NA |

**Table S1.** Taxon, sample locality, number of reads, number of contigs, and number of UCEs for each individual.

| **Taxon** | **Sample Locality** | **Reads** | **Contigs** | **UCEs** |
| --- | --- | --- | --- | --- |
| *Pantherophis obsoletus* | Jefferson LA | 2192806 | 13226 | 4050 |
| *Pituophis catenifer* | Sioux NE | 2138167 | 9525 | 3937 |
| *Pituophis catenifer* | Wheeler TX | 2789601 | 16526 | 3705 |
| *Pituophis ruthveni* | Bienville LA_1 | 1834169 | 8905 | 4000 |
| *Pituophis ruthveni* | Bienville LA_2 | 1940223 | 10197 | 4082 |
| *Pituophis m. lodingi* | Jackson MS_1 | 2476764 | 11792 | 3830 |
| *Pituophis m. lodingi* | Jackson MS_2 | 1595826 | 8416 | 4076 |
| *Pituophis m. lodingi* | Jackson MS_3 | 1591374 | 8514 | 4077 |
| *Pituophis m. lodingi* | Mobile AL | 2100669 | 10270 | 4105 |
| *Pituophis m. lodingi* | Perry MS_1 | 1414642 | 8744 | 3867 |
| *Pituophis m. lodingi* | Perry MS_2 | 1356673 | 7619 | 3889 |
| *Pituophis m. lodingi* | Perry MS_3 | 1097785 | 7715 | 3897 |
| *Pituophis m. melanoleucus* | Autauga AL | 2116151 | 9591 | 3909 |
| *Pituophis m. melanoleucus* | Burlington NJ_1 | 1061120 | 8046 | 3858 |
| *Pituophis m. melanoleucus* | Burlington NJ_2 | 1336710 | 9439 | 3868 |
| *Pituophis m. melanoleucus* | Humphreys TN | 1854737 | 12198 | 3913 |
| *Pituophis m. melanoleucus* | Huntingdon TN | 1630352 | 8578 | 3992 |
| *Pituophis m. melanoleucus* | Marion GA_1 | 1853974 | 12478 | 4011 |
| *Pituophis m. melanoleucus* | Marion GA_2 | 2613221 | 19784 | 3436 |
| *Pituophis m. melanoleucus* | Ocean NJ | 1950980 | 10749 | 3828 |
| *Pituophis m. melanoleucus* | Richmond NC | 1671822 | 9973 | 3969 |
| *Pituophis m. melanoleucus* | Rockingham NC | 1822674 | 9829 | 3919 |
| *Pituophis m. melanoleucus* | Scotland NC | 1824724 | 7731 | 4156 |
| *Pituophis m. melanoleucus* | Winston AL | 2356280 | 14303 | 3810 |
| *Pituophis m. melanoleucus* | Hoke NC | 1335112 | 7503 | 4033 |
| *Pituophis m. mugitus* | Baker GA | 2038991 | 11954 | 4041 |
| *Pituophis m. mugitus* | Covington AL_1 | 2787593 | 13899 | 3910 |
| *Pituophis m. mugitus* | Covington AL_2 | 2628895 | 14689 | 3832 |
| *Pituophis m. mugitus* | Pierce GA | 2422072 | 10629 | 4115 |
| *Pituophis m. mugitus* | Turner GA | 1580711 | 8190 | 4039 |
| *Pituophis m. mugitus* | Alachua FL | 1830880 | 10553 | 4080 |
| *Pituophis m. mugitus* | Bristol FL | 2882788 | 15246 | 3622 |
| *Pituophis m. mugitus* | Brooksville FL | 1618837 | 7840 | 4089 |
| *Pituophis m. mugitus* | Cantonment FL | 3491931 | 14175 | 3832 |
| *Pituophis m. mugitus* | Hamilton FL | 1845695 | 12142 | 3895 |
| *Pituophis m. mugitus* | Hernando FL | 1255625 | 7583 | 3848 |
| *Pituophis m. mugitus* | Highlands FL | 2214725 | 12322 | 4032 |
| *Pituophis m. mugitus* | Lafayette FL | 2314768 | 10400 | 3903 |
| *Pituophis m. mugitus* | Liberty FL | 1526608 | 9735 | 3860 |
| *Pituophis m. mugitus* | Marion FL | 1308565 | 7136 | 4056 |
| *Pituophis m. mugitus* | Okaloosa FL | 1353711 | 8932 | 3709 |
| *Pituophis m. mugitus* | Okeechobee FL_1 | 1722263 | 11758 | 3383 |
| *Pituophis m. mugitus* | Okeechobee FL_2 | 1759949 | 11434 | 3970 |
| *Pituophis m. mugitus* | PonceDe Leon FL | 1202307 | 8690 | 3902 |
| *Pituophis m. mugitus* | Santa Rosa FL | 1128023 | 7291 | 3832 |

**Table S2.** Results from generalized dissimilarity modeling (gdm) including backwards elimination of individual predictor variables. The full model includes all seven bioclim variables determined to lack autocorrelation with other bioclim variables, elevation, and river drainage. Each successive model includes all variables from the previous model barring the predictor variable with the least contribution to percent deviance explained. Significance was determined in gdm through 100 permutations. Note that we highlighted the independent predictor variables determined to be significant in our original full model.

| Predictor variable | Full model |  | Model one |  | Model two |  | Model three |  | Model four |  | Model five |  | Model six |  | Model seven |  | Model eight |  |
| --- | --- | --- | --- | --- | --- | --- | --- | --- | --- | --- | --- | --- | --- | --- | --- | --- | --- | --- |
|  | % Deviance | P value | % Deviance | P value | % Deviance | P value | % Deviance | P value | % Deviance | P value | % Deviance | P value | % Deviance | P value | % Deviance | P value | % Deviance | P value |
| **Geographic distance** | **6.18** | **0** | 6.18 | 0 | 6.18 | 0 | 7.01 | 0 | 7.24 | 0 | 8.72 | 0 | 25.85 | 0 | 63.2 | 0 | 63.47 | 0 |
| Elevation | 0.14 | 0.64 | 0.14 | 0.62 | 0.14 | 0.65 | 0.12 | 0.63 | NA | NA | NA | NA | NA | NA | NA | NA | NA | NA |
| **Mean diurnal range** | **21.42** | **0.01** | 21.42 | 0 | 21.42 | 0.02 | 21.47 | 0.01 | 21.45 | 0.02 | 21.41 | 0.02 | 21.99 | 0.01 | 23.59 | 0.01 | 24.32 | 0.02 |
| Max temp warmest month | 0.92 | 0.45 | 0.92 | 0.48 | 0.92 | 0.44 | 0.9 | 0.35 | 0.84 | 0.4 | 0.8 | 0.44 | NA | NA | NA | NA | NA | NA |
| Mean temp wettest month | 0 | 0.99 | 0 | 1 | NA | NA | NA | NA | NA | NA | NA | NA | NA | NA | NA | NA | NA | NA |
| Mean temp driest quarter | 0.03 | 0.66 | 0.03 | 0.67 | 0.03 | 0.7 | NA | NA | NA | NA | NA | NA | NA | NA | NA | NA | NA | NA |
| Annual precipitation | 0.08 | 0.61 | 0.08 | 0.62 | 0.08 | 0.69 | 0.06 | 0.61 | 0.05 | 0.71 | NA | NA | NA | NA | NA | NA | NA | NA |
| Precipitation of the driest month | 0 | 1 | NA | NA | NA | NA | NA | NA | NA | NA | NA | NA | NA | NA | NA | NA | NA | NA |
| **Precipitation of the warmest quarter** | **10.08** | **0.01** | 10.08 | 0.02 | 10.08 | 0.02 | 10.05 | 0.01 | 11.39 | 0.03 | 11.34 | 0.03 | 11.01 | 0.01 | 10.29 | 0.08 | NA | NA |
| **River drainage** | **2.09** | **0.04** | 2.09 | 0.05 | 2.09 | 0.04 | 2.06 | 0.02 | 2.01 | 0.02 | 2.26 | 0.04 | 1.53 | 0.08 | NA | NA | NA | NA |

**Supplemental Figure 1.**
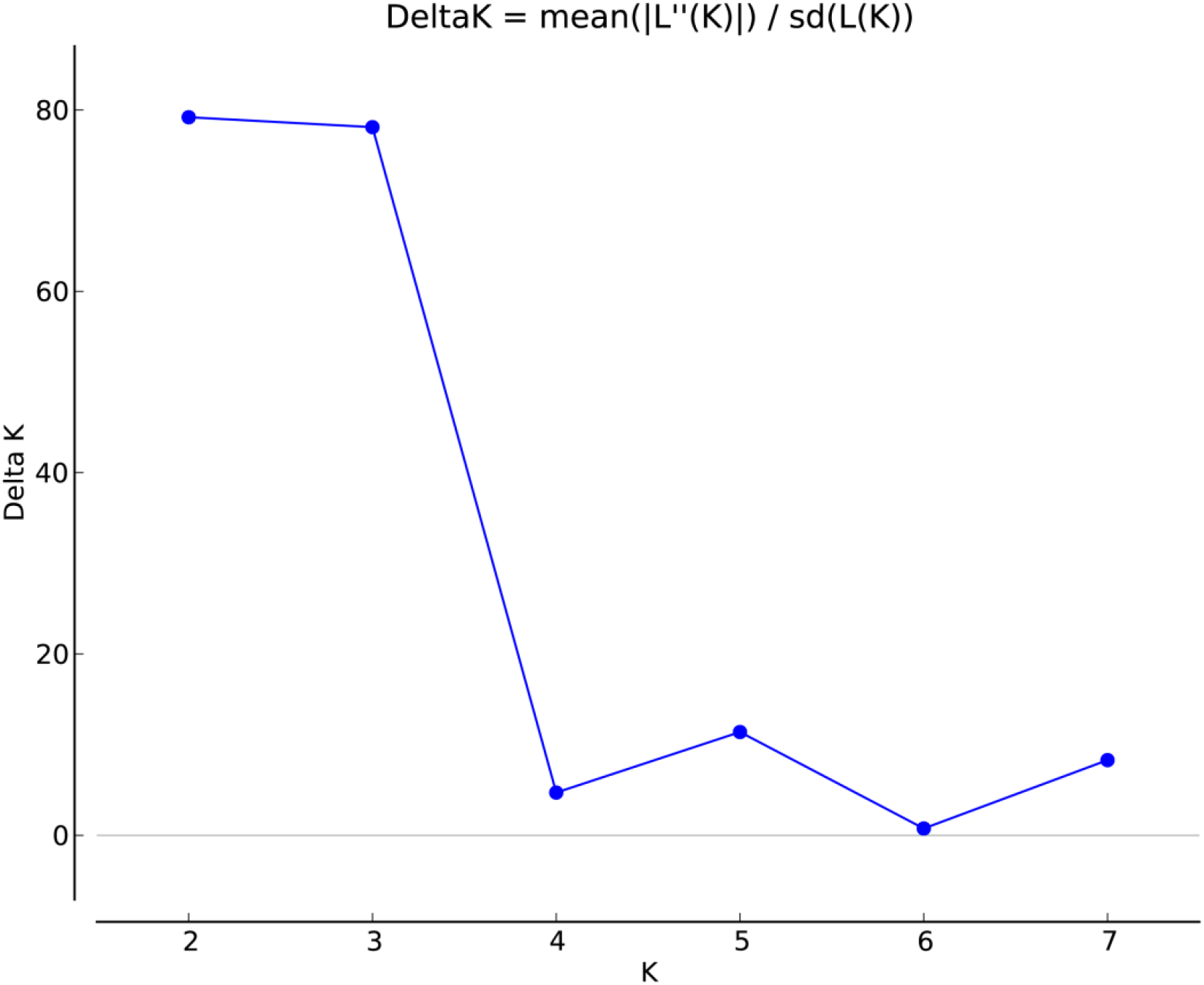
Output from STRUCTURE Harvester. Line plot showing support for a *K*=2 model using the Delta K method for the STRUCTURE analysis.

**Supplemental Figure 2.**
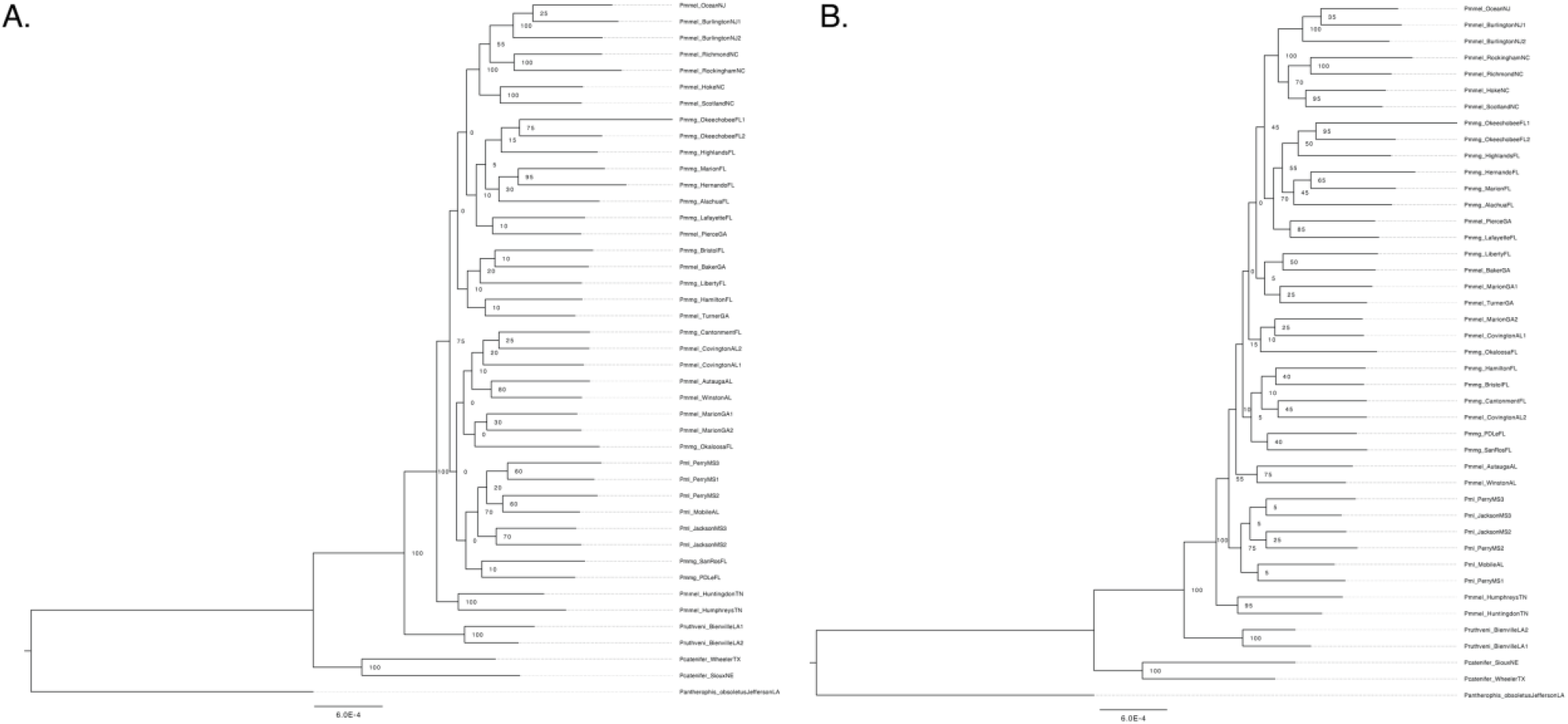
Maximum likelihood tree from A) 50% and B) 75% UCE data matrices.

**Supplemental Figure 3.**
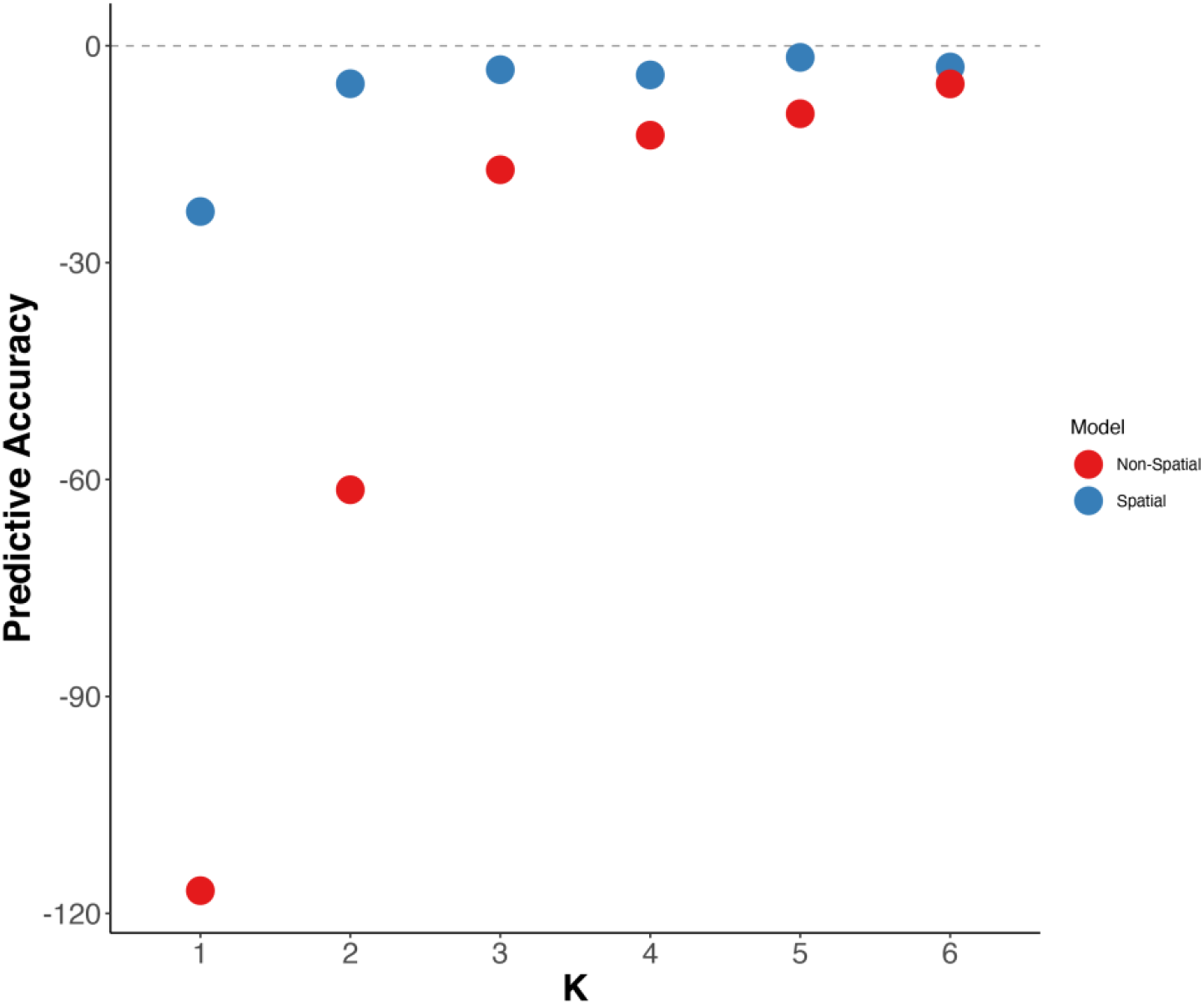
Output from conStruct showing the predictive accuracy for both non-spatial and spatial models from values of *K*=1 to *K*=6.

**Supplemental Figure 4.**
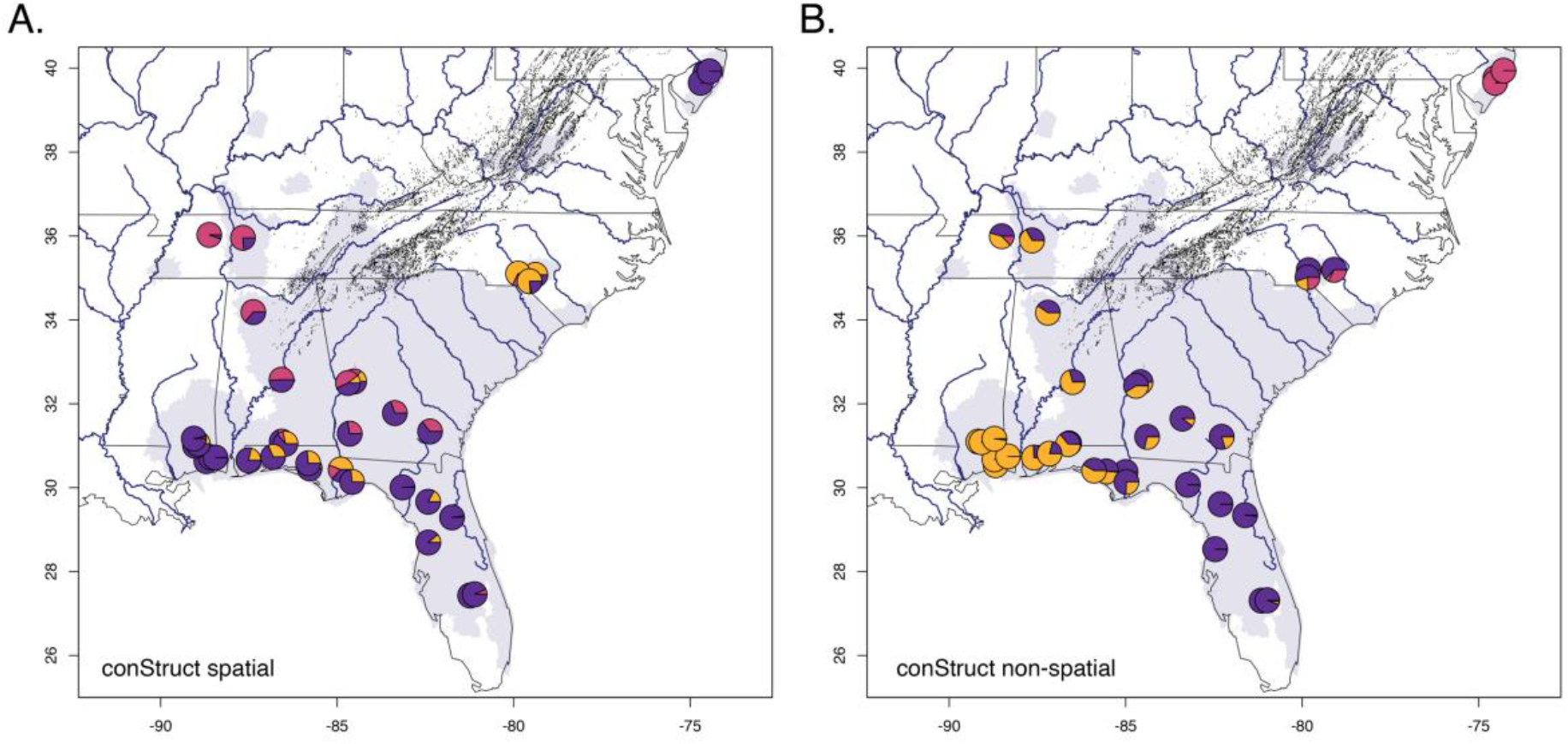
Results from conStruct spatial and non-spatial models. Map of samples with individuals shown as pie charts for *K*=3 model for both spatial (A) and non-spatial (B) models.

## Notes

### Competing Interest Statement

The authors have declared no competing interest.

